# Antibacterial and Anticancer Activities of Insulicolides from *Streptomyces* sp. UL09; An Endophyte of *Ulva rigida* C. Agardh

**DOI:** 10.64898/2026.09.20.751976

**Authors:** Thongchai Taechowisan, Thanaporn Chuen-Im, Waya S. Phutdhawong

## Abstract

Marine macroalgae host endophytic microbial communities that serve as valuable reservoirs for drug discovery. In this study, the endophytic actinomycete *Streptomyces* sp. UL09 was isolated from the internal thallus tissues of the green macroalga *Ulva rigida* C. Agardh. Fermentation of strain UL09 yielded the nitrobenzoyl sesquiterpenoids insulicolide B (1.63 mg/L) and insulicolide C (1.79 mg/L). The planar structures of compounds **1** and **2** were assigned as insulicolides B and C, respectively, via spectroscopic analysis and comparison with literature data. *In vitro* bioassays revealed moderate activity against Gram-positive bacteria (MIC 128–512 µg/mL), including MRSA Sp3 (MIC 128–256 µg/mL). These compounds exhibited cytotoxic activity against human cancer cell lines (HeLa, HepG2, and MDA-MB-231) with IC_50_ values ranging from 34.35 to 142.64 µg/mL. Both compounds demonstrated lower cytotoxicity toward non-malignant Vero cells. Molecular docking simulations predicted that both metabolites target the catalytic ATP-binding cleft of human Polo-like kinase 1 (Plk1), with insulicolide B displaying a favorable docking score (ΔG = -9.273 kcal/mol). Furthermore, pkCSM predicted high intestinal absorption and no alerts for the assessed toxicity endpoints. These findings establish *Streptomyces* sp. UL09 as a viable alternative biological source of insulicolides B and C, highlighting their potential as leads for oncology applications.

## 1. INTRODUCTION

Endophytes encompass a diverse group of bacteria and fungi that reside within the healthy internal tissues of plants during at least one stage of their life cycle, forming symbiotic or mutualistic alliances with their hosts [1]. By occupying this protected internal niche, these microorganisms gain a significant competitive advantage, effectively shielding themselves from the fierce microbial competition typical of the surrounding soil [2]. Among these internal residents, endophytic actinomycetes frequently isolated from rigorously surface-sterilized plant tissues have garnered substantial attention as a highly promising reservoir for novel bioactive natural products [3]. These endophytes synthesize unique metabolic structures that offer unexplored avenues for drug discovery and sustainable agriculture [4].

Recently, marine endophytes have attracted considerable interest as rich, untapped sources of biologically active molecules. Because many soft-bodied marine organisms lack physical defense structures, they rely heavily on chemical defense mechanisms. Survival in these harsh aquatic habitats often depends on the production of bioactive secondary metabolites, synthesized either by the host itself or by its associated microflora [5]. Marine algae, in particular, host diverse bacterial communities that shift dynamically based on the season, host species, and thallus morphology [6,7]. Consequently, actinomycetes associated with marine algae represent a compelling frontier for extensive exploration. Their potential to yield unique, active natural products positions them as an innovative resource for drug discovery that warrants deeper investigation [8].

*Ulva* sp., a genus of green macroalgae within the family Ulvaceae, is well-documented for harboring endophytic bacteria with potent antimicrobial properties [9–12]. A variety of secondary metabolites have been isolated from actinobacteria associated with different *Ulva* species, displaying a broad spectrum of biological activities. For instance, 3-amino-2-carboxamine-6(*R*)-chloro-4(*R*),5(*S*)-dihydroxy-cyclohex-2-en-1-one, 3-amino-2-carboxamine-4(*S*),6(*S*)-dihydroxy-cyclohex-2-en-1-one, and 3-hydroxy-2-propionamidobenzamide were successfully purified from *Streptomyces* sp. ZZ502, an endophyte of *Ulva conglobatea* [13]. Similarly, desertomycin G was obtained from *Streptomyces althioticus* MSM3, an endophyte of *Ulva* sp. [14], while streptopertusacin A and bafilomycins were discovered from *Streptomyces* sp. HZP-2216E, isolated from *Ulva pertusa* [15,16].

Driven by these insights, the present study sought to isolate endophytic actinomycetes from the green macroalga *Ulva rigida* C. Agardh, collected from Bangkaew Beach in Phetchaburi Province, Thailand, to evaluate their efficacy against microbial pathogens. Based on the unique ecological niche of *U. rigida* in this environment, we hypothesize that its endophytic actinomycetes yield novel secondary metabolites featuring potent antibacterial and selective anticancer activities. This investigation focuses on isolating these endophytes, identifying the most potent strains, characterizing their bioactive secondary metabolites, and evaluating their therapeutic potential for oncology and infectious disease applications.

## 2. MATERIALS AND METHODS

### 2.1. Sample Collection and Endophytic Isolation

Three healthy specimens of the green macroalga *Ulva rigida* C. Agardh were collected from Bangkaew Beach, Banlaem District, Phetchaburi Province, Thailand (13.109583°N, 100.060722°E). For endophyte isolation, the thalli were dissected into small segments (approximately 5 x 5 mm^2^), thoroughly rinsed, and subjected to a multi-step surface sterilization protocol as previously described [10]. To ensure the efficacy of the surface sterilization, 100 µL of the final double-distilled wash water was spread onto ISP-2 medium and incubated at 32°C for 7 days. The absence of microbial growth on the culture medium confirmed the effectiveness of the surface sterilization. The sterilized tissue pieces were dried under aseptic conditions in a laminar flow cabinet (Esco Scientific, PA, USA) before being plated onto humic acid-vitamins (HV) agar [17]. The medium was supplemented with 100 µg/mL each of cycloheximide and nystatin to suppress fungal and yeast overgrowth.

Following a 3-week incubation period at 32°C, colonies displaying distinct actinomycete morphology were isolated and subcultured onto fresh International *Streptomyces* Project medium 2 (ISP-2) agar plates [18] to obtain pure cultures. A total of 38 actinomycete isolates were initially screened for antibacterial efficacy using the soft-agar overlay technique [19]. The strain designated UL09 exhibited the most prominent zone of inhibition against the panel of indicator bacteria and was selected for comprehensive taxonomic identification and downstream analysis.

### 2.2. Taxonomic Identification of Strain UL09

Morphological attributes of the selected strain were analyzed via scanning electron microscopy using a TESCAN Mira3 system (TESCAN, Czech Republic). Chemotaxonomic profiling and 16S rRNA gene sequencing were executed according to the methods established by Taechowisan *et al*. [20].

### 2.3. Fermentation and Metabolite Extraction

For large-scale metabolite production, strain UL09 was inoculated onto 500 Petri dishes containing ISP-2 agar (approximately 10 L total volume) and incubated at 32 °C for 21 days. The resulting agar cultures were thoroughly extracted with ethyl acetate (EtOAc) to recover secondary metabolites [20]. The organic phases were pooled and concentrated under reduced pressure using a rotary evaporator, yielding approximately 18.42 g of a dark brown crude extract. This crude material was partitioned into two aliquots: one fraction was dissolved in dimethyl sulfoxide (DMSO) for in vitro bioassays, while the remainder was solubilized in dichloromethane (CH_2_Cl_2_) and reserved for chromatographic separation.

### 2.4. Purification and Structural Characterization

A portion of the crude extract (∼15.00 g) was fractionated using silica gel column chromatography, employing a step-gradient elution system of EtOAc in CH_2_Cl_2_. Fractions showing target compounds that eluted within the 30–35% EtOAc in CH_2_Cl_2_ range were pooled and further purified by preparative thin-layer chromatography (TLC) using an EtOAc:CH_2_Cl_2_ (2:3, v/v) mobile phase. This process yielded purified compound **1** (∼13.27 mg) and compound **2** (∼14.56 mg).

The chemical frameworks of both pure compounds were elucidated using comprehensive spectroscopic techniques. Melting points were recorded, and ultraviolet (UV) absorption spectra were acquired. Detailed electronic and atomic structural profiles were obtained via ^1^H-NMR (500 MHz) and ^13^C-NMR (125 MHz) spectroscopy on a Bruker Avance-500 NMR spectrometer (Bruker, Germany). High-resolution molecular weights were determined using an LTQ Orbitrap XL mass spectrometer (Thermo Fisher Scientific, USA).

### 2.5. Determination of MIC and MBC

Minimum inhibitory concentration (MIC) and minimum bactericidal concentration (MBC) values were determined using a modified broth microdilution method based on Miller *et al*. [21]. Test compounds were dissolved in dimethyl sulfoxide (DMSO) and serially diluted twofold (0.5 − 512 μg/mL) in 96-well microplates containing Brain Heart Infusion (BHI) broth, followed by inoculation with 10 μL of bacterial suspension (10^8^ cells/mL). Chloramphenicol served as the positive control, and broth containing 5% (v/v) DMSO without test compounds was used as the growth control. Plates were incubated at 37°C for 24 h. The MIC was defined as the lowest concentration preventing visible turbidity. To measure MBC, 50 μL from each clear MIC well was plated onto BHI agar and incubated at 37°C for 24 h. The MBC was recorded as the lowest concentration yielding no colony growth. All assays were conducted in triplicate.

### 2.6. *In vitro* Cytotoxicity and Anticancer Assays

The antiproliferative potential of the crude extract and purified metabolites was determined using the standard MTT assay against three human cancer cell lines: HeLa (cervical carcinoma), HepG2 (hepatocellular carcinoma), and MDA-MB-231 (breast adenocarcinoma) [22]. General cytotoxicity evaluations were performed in parallel on the non-cancerous Vero cell line. All samples were assayed across a concentration gradient spanning 1-512 µg/mL. The Selectivity Index (SI) was determined using the following equation:

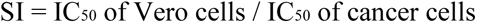

Doxorubicin hydrochloride (Thermo Fisher Scientific, USA) served as the positive control.

### 2.7. Molecular Docking Simulations

Three-dimensional (3D) structural models of the target nitrobenzoyl sesquiterpenoids, insulicolide B and insulicolide C, were generated and energy-minimized utilizing UCSF Chimera software [23]. The 3D crystal structure of human Polo-like kinase 1 (Plk1), an established cell-cycle regulatory target for oncology therapeutics [24], was retrieved from the Protein Data Bank (PDB ID: 2OWB). Docking simulations were performed via AutoDock Vina within the UCSF Chimera interface [25]. A 20 x 20 x 20 Å cubic grid box centered over the active site was defined based on the coordinates of the co-crystallized native inhibitor, 4-(4-methylpiperazin-1-yl)-N-[5-(2-thiophen-2-ylethanoyl)-1H-pyrrolo[3,4-c]pyrazol-3-yl]benzamide (compound 626; PubChem ID: 11963557, DrugBank ID: DB07186). Binding affinities were calculated in kcal/mol, and the lowest-energy conformations were identified as the optimal docking orientations. Non-covalent intermolecular variations (hydrophobic contacts and hydrogen bonding) were mapped using Discovery Studio Visualizer (BIOVIA). Doxorubicin was included as the reference comparative ligand.

### 2.8. *In silico* ADMET Predictions

The pharmacokinetic profile encompassing Absorption, Distribution, Metabolism, Excretion, and Toxicity (ADMET) features of the isolated ligands was computed using the online prediction platform pkCSM [26]. Doxorubicin was analyzed concurrently under identical parameters to serve as a baseline reference.

### 2.9. Statistical Analysis

All biological evaluations were conducted in triplicate using independent experimental runs. Quantitative values are expressed as the mean ± standard deviation (SD). Statistical significance between groups was determined via one-way Analysis of Variance (ANOVA) followed by Tukey’s post hoc test, implemented using SPSS for Windows, version 11.01 (SPSS Inc., Chicago, IL, USA). The threshold for statistical significance was defined at *p* < 0.05.

## 3. RESULTS

The processing of *Ulva rigida* C. Agardh thalli successfully yielded culturable endophytic actinomycetes [Figure 1a]. Following a 3-week incubation period on HV agar, distinct dry, powdery, white actinomycete colonies emerged directly from the margins of the inoculated tissue segments [Figure 1b].

**Figure 1:**
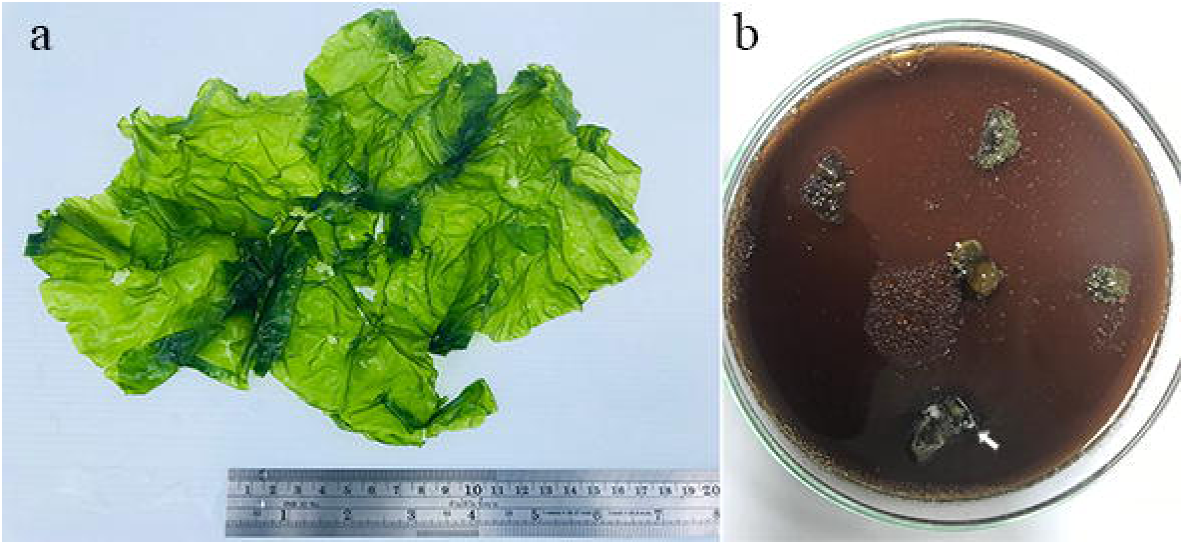
Isolation workflow and macroalgal source material. (a) Morphological appearance of the *Ulva rigida* C. Agardh thallus collected from Bangkaew Beach, Phetchaburi Province, Thailand. (b) Emergence of distinct endophytic actinomycete colonies (marked by white arrows) radiating from the margins of surface-sterilized *U. rigida* tissue segments after a 3-week cultivation period on humic acid-vitamins (HV) agar at 32°C.

From 150 tissue segments obtained across three independent *U. rigida* thallus samples, 38 distinct isolates were recovered, representing an isolate recovery rate of 25.3% per segment. Initial antimicrobial screening via the soft-agar overlay method identified a single candidate, designated as strain UL09, which exhibited strong antagonistic activity. This specific isolate generated prominent zones of inhibition ranging from 37 mm to 43 mm against the target pathogens [Figure 2]. On the basis of this robust bioactivity, strain UL09 was prioritized for comprehensive taxonomic characterization and molecular identification.

**Figure 2:**
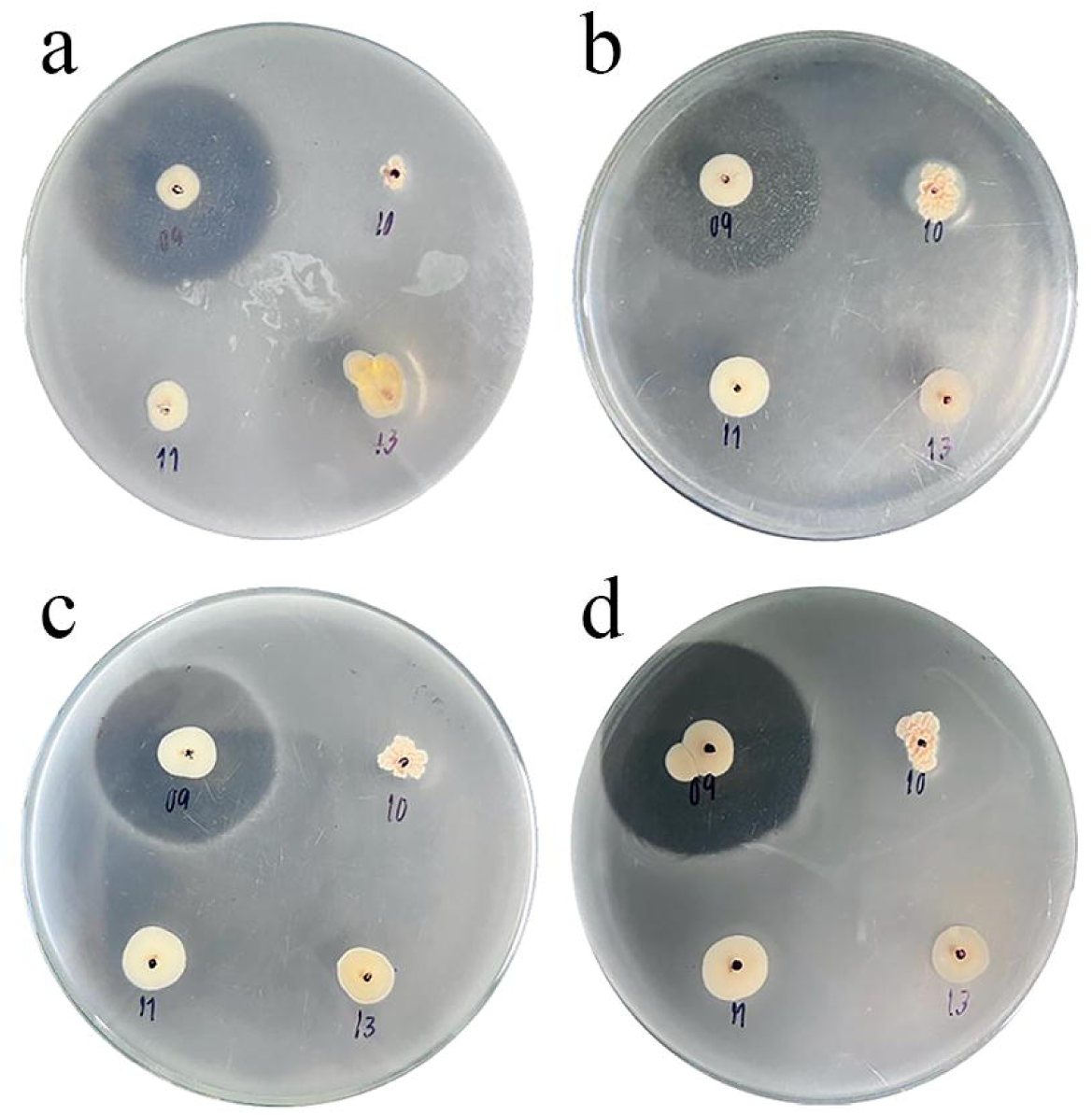
Screening of strain UL09 for antimicrobial efficacy using the soft-agar overlay method. Zones of growth inhibition were evaluated against target pathogens after overlaying 7-day-old *Streptomyces* sp. UL09 colonies on ISP-2 agar and incubating for 24 hours at 37°C. Representative bioactivity panels illustrate antagonistic performance against (a) *Bacillus cereus* TISTR687, (b) *Escherichia coli* TISTR887, (c) methicillin-resistant *Staphylococcus aureus* (MRSA) Sp3, and (d) *Staphylococcus epidermidis* TISTR518.

Cultivation of isolate UL09 on ISP-2 agar yielded characteristic leathery colonies featuring a distinct pale sandstone coloration [Figure 3a]. The young aerial mycelia were initially white, transitioning into a rich cappuccino hue after 7 days of incubation. Morphological examination via scanning electron microscopy (SEM) revealed well-developed, non-fragmenting substrate and aerial mycelia, with the latter organized into monopodially branched sporophores. The resulting spores were flexible, rod-shaped, and characterized by a smooth surface coat [Figure 3b].

**Figure 3:**
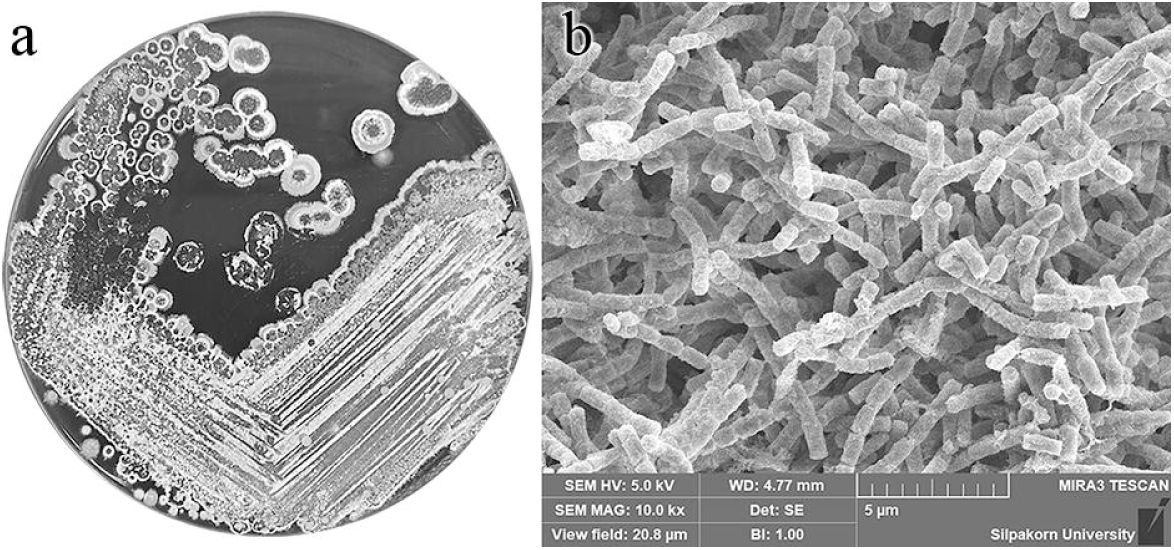
Morphological characteristics of *Streptomyces* sp. UL09. (a) Macromorphology of strain UL09 displaying characteristic vegetative growth on ISP-2 agar following a 15-day incubation period at 32°C. (b) Scanning electron micrograph (SEM) showing the ultrastructure of the aerial mycelium, featuring flexible spore chains composed of rod-shaped spores with a smooth surface texture.

Chemotaxonomic evaluation of the whole-cell wall hydrolysates revealed the diagnostic presence of *LL*-diaminopimelic acid (*LL*-DAP). The detection of this specific isomer confirmed the alignment of the isolate with the cell wall chemotype characteristic of the genus *Streptomyces*.

Taxonomic assignment was resolved by sequencing the 16S rRNA gene of strain UL09. A localized BLAST search of the assembled sequence demonstrated a high degree of sequence homology, sharing 98.71% identity with the type strain *Streptomyces qinzhouensis* SSL-25^T^. Phylogenetic reconstruction utilizing the Neighbor-Joining method corroborated this relationship, positioning strain UL09 within a robust, tightly clustered clade alongside *S. qinzhouensis* [Figure 4]. The complete 16S rRNA gene sequence for strain UL09 has been formally deposited in the GenBank database under the accession number PZ626752.

**Figure 4:**
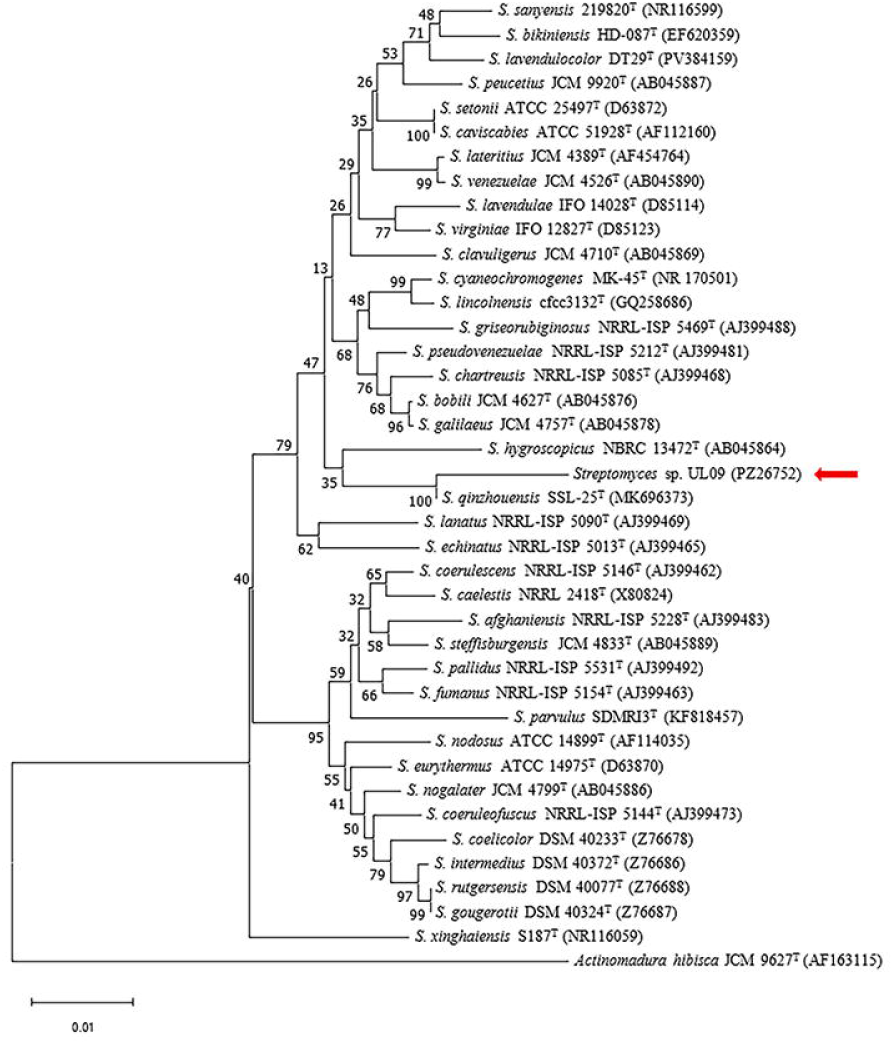
Evolutionary relationships of strain UL09. Phylogenetic reconstruction based on 16S rRNA gene sequences showing the position of *Streptomyces* sp. UL09 relative to closely related *Streptomyces* type strains retrieved from GenBank (accession numbers are indicated in parentheses). The tree was generated via the Neighbor-Joining method using MEGA11 software. Topology confidence was assessed through bootstrap analysis with 1,000 replicates, and percentages are shown at the nodes. The scale bar corresponds to 0.01 substitutions per nucleotide site.

Scale-up fermentation of *Streptomyces* sp. UL09 (10 L) yielded 18.42 g of total crude extract. A 15.00 g aliquot was subjected to sequential silica gel column chromatography, yielding 13.27 mg of compound **1** and 14.56 mg of compound **2**. To reflect total batch production, the final yield values (1.63 mg/L for compound **1** and 1.79 mg/L for compound **2**) were calculated by extrapolating these recovered masses to the full 18.42 g crude extract and dividing by the culture volume.

Compound **1**: Obtained as a white amorphous powder (MP: 184–186°C; yield: 1.63 mg/L of culture medium), MP 184°C−186°C, UV (MeOH)λ_max_ nm (log*ε*): 201.7 (0.028), 254.2 (0.021), 292.3 (0.006), 369.5 (0.004); IR*υ*_max_ (KBr) cm^-1^: 3385, 3024, 2967, 2832, 1713, 1642, 1556, 1250, 1164, 1107, 1060; HRESIMS m/z (rel. int.): 416.1706 [M + H] ^+^ (calcd. for C_22_H_25_NO_7_, 416.1704; mass error: +0.4806 ppm). The ^1^H-NMR data (CD_3_OD, δ, ppm, *J*/Hz): 8.36 (2H, brd, *J* = 8.9, H-4’, H-6’), 8.24 (2H, brd, *J* = 8.8, H-3’, H-7’), 6.77 (1H, dd, *J* = 3.6, 3.6, H-7), 4.98 − 5.13 (2H, d, *J* = 11.4, 11.4, H-14), 4.76 (1H, dd, *J* = 4.8, 3.6, H-6), 4.18 − 4.51 (2H, dd, *J* = 9.2, 9.2, H-11), 2.87 (1H, m, H-9), 1.19 − 2.09 (2H, ddd, *J* = 13.4, 13.4, 2.9, brd, *J* = 13.4, H-3), 1.56 (1H, d, *J* = 4.8, H-5), 1.50 − 1.72 (2H, m, H-2), 1.37 − 1.72 (2H, ddd, *J* = 13.7, 13.7, 3.9, brd, *J* = 13.7, H-1), 1.33 (3H, s, H-13), 1.09 (3H, s,H-15). ^13^C-NMR data (CD_3_OD, δ, ppm, *J*/Hz): 171.8 (s, C-12), 165.5 (s, C-1’), 151.4 (s, C-5’), 136.8 (d, C-7), 136.4 (s, C-2’), 130.9 (d, C-3’, C-7’), 127.8 (s, C-8), 124.0 (d, C-4’, C-6’), 69.2 (t, C-14), 68.6 (t, C-11), 64.4 (d, C-6), 57.0 (d, C-5), 52.0 (d, C-9), 41.0 (t, C-1), 38.8 (s, C-4), 37.1 (t, C-3), 34.6 (s, C-10), 26.3 (q, C-13), 18.3 (t, C-2), 15.6 (q, C15).

Compound **2**: Obtained as a white amorphous powder (MP: 185–187°C; yield: 1.79 mg/L of culture medium), UV (MeOH)λ_max_ nm (log*ε*): 200.9 (0.029), 253.1 (0.020), 290.8 (0.007), 368.7 (0.005); IR*υ*_max_ (KBr) cm^-1^: 3387, 3020, 2972, 2836, 1721, 1653, 1552, 1258, 1167, 1109, 1056; HRESIMS m/z (rel. int.): 458.1813 [M + H] ^+^ (calcd. for C_24_H_27_NO_8_, 458.1810; mass error: +0.6548 ppm). The ^1^H-NMR data (CD_3_OD, δ, ppm, *J*/Hz): 8.36 (2H, brd, *J* = 9.0, H-4’, H-6’), 8.25 (2H, brd, *J* = 9.0, H-3’, H-7’), 6.75 (1H, dd, *J* = 3.8, 3.8, H-7), 6.15 (1H, m, H-6), 4.25 − 4.62 (2H, d, *J* = 11.3, 11.3, H-14), 4.24 − 4.57 (2H, dd, *J* = 9.2, 9.2, H-11), 3.04 (1H, m, H-9), 1.95 (1H, d, *J* = 4.3, H-5), 1.91 (3H, s, H-2”), 1.53 − 1.68 (2H, m, H-2), 1.48 − 2.05 (2H, ddd, *J* = 13.5, 13.5, 3.9, brd, *J* = 13.5, H-1), 1.26 (3H, s, H-15), 1.21 − 1.81 (2H, m, brd, *J* = 13.3, H-3), 1.12 (3H, s, H-13). ^13^C-NMR data (CD_3_OD, δ, ppm, *J*/Hz): 172.1 (s, C-1”), 170.7 (s, C-12), 164.3 (s, C1’), 151.6 (s, C-5’), 135.5 (s, C-2’), 132.2 (s, C-8), 131.4 (d, C-3’, C-7’), 129.8 (d, C-7), 124.1 (d, C-4’, C-6’), 68.6 (d, C-6), 68.4 (t, C-11), 67.4 (t, C-14), 54.9 (d, C-5), 52.0 (d, C-9), 40.9 (t, C-1), 38.1 (s, C-4), 37.3 (t, C-3), 35.1 (s, C-10), 26.1 (q, C-13), 19.8 (q, C-2”), 18.1 (t, C-2), 16.1 (q, C-15).

Based on UV, ^1^H-NMR, ^13^C-NMR, and HRESIMS data, compounds **1** and **2** were assigned as insulicolide B and insulicolide C, respectively, by comparison with published planar spectra [Figure 5]. Both metabolites belong to the sesquiterpenoid chemical class.

**Figure 5:**
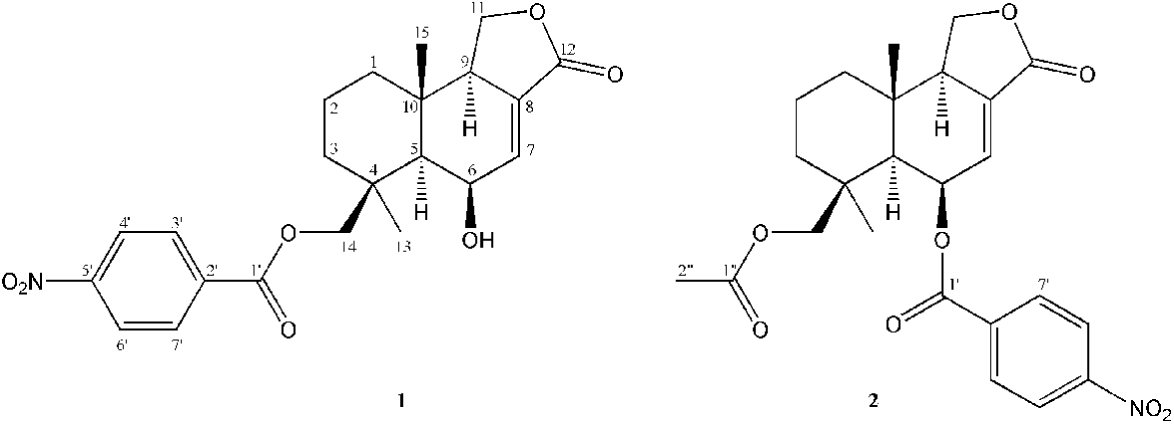
Structures of the compounds. [(5*R*,5*aR*,6*S*,9*aR*,9*bR*)-5-hydroxy-6,9*a*-dimethyl-3-oxo-5,5*a*,7,8,9,9*b*-hexahydro-1*H*-benzo[e][2]benzofuran-6-yl]methyl 4-nitrobenzoate (**1**) and [(5*R*,5*aR*,6*S*,9*aR*,9*bR*)-6-(acetyloxymethyl)-6,9*a*-dimethyl-3-oxo-5,5*a*,7,8,9,9*b*-hexahydro-1*H*-benzo[e][2]benzofuran-5-yl] 4-nitrobenzoate (**2**).

The purified sesquiterpenoids demonstrated moderate antibacterial activity, primarily targeting Gram-positive bacterial strains. Against sensitive targets, including the clinical methicillin-resistant *Staphylococcus aureus* (MRSA) strain Sp3, compound MIC values ranged from 128 to 512 µg/mL, with corresponding MBC values spanning 512 to greater than 512 µg/mL [Table 1]. Conversely, efficacy against Gram-negative pathogens was poor; both MIC and MBC thresholds reached the maximum evaluated range of 512 to greater than 512 µg/mL.

The antiproliferative profiles of the crude extract and the purified sesquiterpenoids were determined against three human cancer cell lines (HeLa, HepG2, and MDA-MB-231) alongside a non-malignant Vero cell control. Both isolated compounds displayed measurable, dose-dependent efficacy against the malignant cell lines, yielding IC_50_ values between 34.35 and 142.64 µg/mL. Notably, toxicity toward normal Vero cells was low, as indicated by elevated IC_50_ parameters ranging from 234.84 to 323.75 µg/mL.

Against the HepG2 cell line, the Selectivity Index (SI) values of the isolated sesquiterpenoids and the crude extract exceeded the corresponding margin observed for doxorubicin hydrochloride [Table 2]. In contrast, the SI values recorded for the MDA-MB-231 and HeLa cell lines were lower than those calculated for doxorubicin. Taken together, these data establish *U. rigida* thallus tissue as a viable source for isolating *Streptomyces* sp. UL09, a strain capable of producing sesquiterpenoid metabolites with selective antiproliferative activity.

To investigate the putative molecular mechanisms governing the observed anticancer profiles, molecular docking simulations were conducted to evaluate the interactions of compounds **1** and **2** within the catalytic domain of human Polo-like kinase 1 (Plk1; PDB ID: 2OWB). Doxorubicin and the native inhibitor compound 626 were modeled in parallel as reference standards.

The calculated relative binding energies (ΔG) were −9.273 kcal/mol for compound **1**, −8.641 kcal/mol for compound **2**, −9.112 kcal/mol for doxorubicin, and −8.938 kcal/mol for compound 626. All four test ligands stabilized within the active pocket of the Plk1 catalytic domain, driven primarily by an interplay of hydrophobic forces and targeted hydrogen bonds [Figure 6].

**Figure 6:**
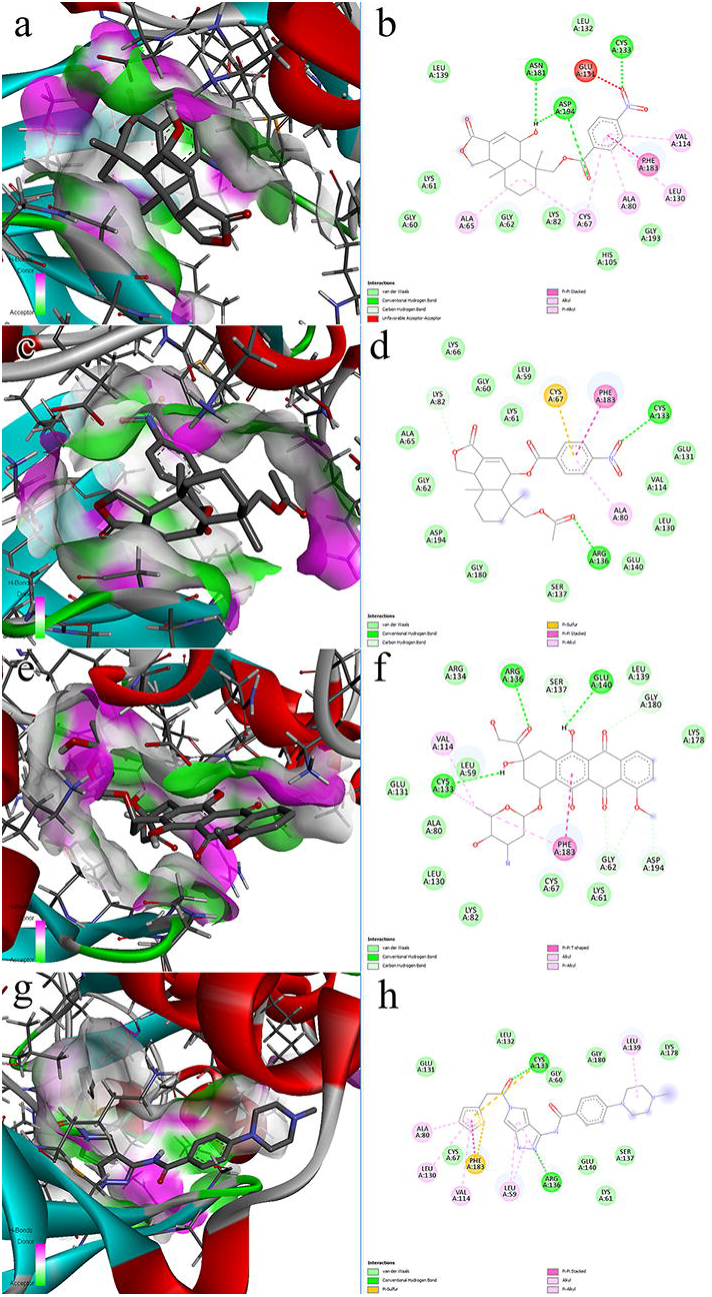
Molecular docking profiles within the active site of human Polo-like kinase 1 (Plk1; PDB ID: 2OWB). The panels illustrate the three-dimensional (3D) binding conformations and matching two-dimensional (2D) interaction networks for: (a, b) compound **1**, (c, d) compound **2**, (e, f) doxorubicin, and (g, h) reference compound 626. In the 3D models, hydrogen bond donor and acceptor atoms within the binding pocket are color-coded in pink and green, respectively. In the 2D schematics, non-covalent interactions are designated as follows: conventional hydrogen bonds (dark green dashes), carbon-hydrogen bonds (light green dashes), Pi-Pi-stacked and Pi-Pi-T-shaped interactions (deep pink dashes), Pi-alkyl and alkyl interactions (pale pink dashes), Pi-sulfur interaction (pale brown dashes). Unfavorable acceptor-acceptor interaction is indicated by red dashes, while residues participating in van der Waals contacts are shaded in pale green.

Specifically, compound **1** established hydrogen bonding networks with Cys133, Asn181, and Asp194. Compound **2** similarly formed an H-bond with Cys133, but was uniquely stabilized by additional H-bond anchors to Arg136 and Lys82. The reference molecules (doxorubicin and compound 626) also shared the H-bond linkage with Arg136 [Table 3]. Compound **1** exhibited a favorable docking score (ΔG = −9.273 kcal/mol), comparable to doxorubicin (−9.112 kcal/mol) and compound 626 (−8.938 kcal/mol). Although compound **2** yielded a slightly less negative binding energy (−8.641 kcal/mol), the docking simulations suggest that both isolated sesquiterpenoids adopt favorable predicted binding poses within the Plk1 catalytic site. Detailed non-covalent mapping, including supplementary van der Waals contacts with residues Leu59, Gly60, Lys61, Gly62, Ala65, Lys66, Lys82, His105, Val114, Leu130, Gly131, Leu132, Ser137, Leu139, Glu140, Gly180, Gly193, and Asp194, is compiled in Figure 6.

Predictive ADMET parameters for the isolated ligands were generated and benchmarked against doxorubicin and compound 626 [Table 4]. Absorption: Both compound **1** (91.93%) and compound **2** (99.79%) exhibited high predicted Human Intestinal Absorption (HIA), distinctly outperforming doxorubicin (62.37%). The sesquiterpenoids also demonstrated elevated Caco-2 monolayer permeability scores. Neither ligand was predicted to act as a P-glycoprotein (P-gp) substrate, though both were categorized as potential P-gp inhibitors. Distribution: Compounds **1** and **2** showed predicted blood-brain barrier penetration (logBB) values of -0.743 and -1.085, with corresponding central nervous system accumulation (logPS) values of -2.452 and -2.818. Metabolism: The isolated sesquiterpenoids were predicted to act as substrates and inhibitors of the CYP3A4 isoform, but they did not show substrate or inhibitory activity toward other major cytochrome P450 enzymes. Excretion: Neither compound was flagged as an organic cation transporter 2 (OCT2) substrate. The predicted total clearance values for compounds **1** and **2** were lower than those of doxorubicin and compound 626. Toxicity: Computational toxicological screening indicated a favorable safety profile; neither compound presented predictive liabilities for Ames mutagenicity, cardiotoxicity, hepatotoxicity, or skin sensitization.

## 4. DISCUSSION

The present study details the isolation and structural elucidation of two sesquiterpenoid derivatives from the endophytic actinomycete *Streptomyces* sp. UL09, recovered from the macroalga *Ulva rigida*. These compounds were resolved as [(5*R*,5*aR*,6*S*,9*aR*,9*bR*)-5-hydroxy-6,9*a*-dimethyl-3-oxo-5,5*a*,7,8,9,9*b*-hexahydro-1*H*-benzo[e][2]benzofuran-6-yl]methyl 4-nitrobenzoate (compound **1**) and [(5*R*,5*aR*,6*S*,9*aR*,9*bR*)-6-(acetyloxymethyl)-6,9*a*-dimethyl-3-oxo-5,5*a*,7,8,9,9*b*-hexahydro-1*H*-benzo[e][2]benzofuran-5-yl] 4-nitrobenzoate (compound **2**). Based on 1D NMR (^1^H and ^13^C) and HRESIMS comparative structural analysis, compounds **1** and **2** show planar identity to insulicolide B and insulicolide C, previously isolated from the endophytic fungus *Aspergillus ochraceus* Jcma1F17 [27]. However, full 2D NMR spectroscopic analysis (COSY, HSQC, HMBC, and NOESY) combined with optical rotation or ECD measurements is required to definitively confirm their relative and absolute stereochemical assignments.

While sesquiterpenoids are distributed across diverse biological taxa including terrestrial plants, fungi, and bacteria, and are noted for broad functional activities [28], their recovery from an algal bacterial endophyte emphasizes the cross-kingdom occurrence of these secondary metabolites. The successful isolation of these compounds from *Streptomyces* sp. UL09 reinforces the value of marine macroalgal endophytes as reservoirs for discovering novel structures or establishing alternative biological sources for known bioactive leads.

Notably, *Streptomyces* sp. UL09 produced quantifiable amounts of both compounds under standard ISP-2 culture conditions without prior media optimization. Strain UL09 yielded 1.63 mg/L of compound **1** and 1.79 mg/L of compound **2**. This production was higher than the yields reported from the marine fungal source *A. ochraceus* Jcma1F17, which produced 0.11 mg/L of insulicolide B and 0.09 mg/L of insulicolide C [27]. This marked difference suggests that *Streptomyces* sp. UL09 possesses an efficient metabolic framework for synthesizing these complex sesquiterpenoids. Consequently, targeted optimization of fermentation parameters such as carbon/nitrogen ratios, temperature profiles, and aeration rates presents a viable pathway for scaling up these production metrics.

Phylogenetic classification clustered strain UL09 with *Streptomyces qinzhouensis*, showing 98.71% 16S rRNA gene sequence similarity to the type strain *S. qinzhouensis* SSL-25^T^ [29].

Although the original type strain was isolated from mangrove soil in Qinzhou Bay, China, its secondary metabolome and biological activities remained uncharacterized [29]. The current findings extend the known environmental scope of this *Streptomyces* lineage to marine macroalgae and link the taxon to bioactive sesquiterpenoid production.

In bioactivity assays, the purified sesquiterpenoids demonstrated moderate antibacterial efficacy against Gram-positive pathogens (MIC = 128 to 512 µg/mL), whereas activity against Gram-negative species was low (MIC ≥ 512 µg/mL). This profile is generally consistent with previously reported *Streptomyces fulvorobeus* sesquiterpenoids, which exhibited weak antibacterial thresholds against select Gram-positive and Gram-negative strains [30].

Sesquiterpenoids frequently exhibit antiproliferative properties through the suppression of cell growth or the induction of apoptotic pathways [31]. In this study, compounds **1** and **2** showed cytotoxic activity against MDA-MB-231, HeLa, and HepG2 cell lines, yielding IC_50_ values between 34.35 and 142.64 µg/mL. From a therapeutic standpoint, evaluating safety margins relative to non-malignant cells is crucial. While the compounds displayed measurable baseline toxicity toward non-cancerous Vero cells (IC_50_ = 234.84 to 323.75 μg/mL), the calculated ratio for the HepG2 liver cancer line relative to Vero cells was higher than that of the reference drug, doxorubicin. This suggests a relatively favorable *in vitro* safety window toward hepatocellular carcinoma cells, mirroring the cytotoxicity patterns reported for sesquiterpenoids derived from *Streptomyces qinglanensis*, which inhibited solid tumor lines with GI_50_ values spanning 1.97 − 3.46 μM [32].

To explore potential binding interactions underlying this antiproliferative effect, molecular docking simulations were performed against human Polo-like kinase 1 (Plk1, PDB ID: 2OWB), a serine/threonine kinase that regulates cell-cycle progression and cytokinesis [24]. Both insulicolide derivatives, along with doxorubicin and the synthetic inhibitor compound 626, accommodated the Plk1 active site with favorable binding energies. Compound **1** exhibited the highest predicted thermodynamic stability (ΔG = −9.273 kcal/mol), outperforming doxorubicin (−9.112 kcal/mol) and compound 626 (−8.938 kcal/mol). Conversely, compound **2** displayed a slightly lower affinity (−8.641 kcal/mol). This study relies on computational predictions suggesting that the purified compounds act as human Polo-like kinase 1 inhibitors. However, this mechanism requires experimental validation and warrants further investigation. In addition, molecular dynamics simulations should be performed for all ligands and reference compounds to further evaluate the stability and behavior of their complexes with human Polo-like kinase 1.

Structural validation of the Plk1 active site using the X-ray crystal structure complexed with the pyrrolo-pyrazole inhibitor PHA-680626 has previously established key binding pocket interactions involving Leu59, Cys67, Ala80, Lys82, Val114, Leu130, Glu131, Cys133, Arg136, Phe183, and Asp194 [33,34]. Furthermore, structural mapping of the anticancer sesquiterpene lactone racemolactone I indicated active site occupancy through interactions with Leu59, Gly60, Lys61, Gly62, Ala65, Lys66, Cys67, Ala80, Lys82, Cys133, Arg136, Ser137, Leu139, Glu140, Lys143, Lys178, Gly180, Asn181, and Phe183 [35].

Our docking data confirm that insulicolides B and C occupy this same catalytic region, forming stable complexes stabilized by overlapping residue networks. A comparative analysis of the binding configurations reveals that Leu59, Gly60, Lys61, Gly62, Ala65, Lys66, Cys67, Ala80, Lys82, Val114, Leu130, Gly131, Leu132, Cys133, Arg136, Ser137, Leu139, Glu140, Gly180, Asn181, Phe183, Gly193, and Asp194 constitute a highly conserved interaction fingerprint shared among the insulicolides, racemolactone I, and PHA-680626. Within this pocket, compound **1** engaged essential ATP-binding residues via hydrogen bonds or tight contacts (His105, Leu132, Leu139, Asn181, and Gly193). In contrast, compound **2** exhibited a distinct stabilization profile, establishing key contacts with Leu59, Lys66, Arg136, Ser137, Glu140, and Gly180. These data support the hypothesis that both sesquiterpenoids act as competitive inhibitors within the ATP-binding cleft of Plk1.

*In silico* ADMET profiling serves as a critical filter for evaluating early-stage drug candidates to manage downstream development risks [26]. The computed pharmacokinetic models predicted that both sesquiterpenoids possess promising drug-like properties in computational models. The predicted Human Intestinal Absorption (HIA) values for compound **1** (91.93%) and compound **2** (99.79%) were superior to the modeled absorption profile of doxorubicin (62.37%), implying a distinct advantage for potential oral delivery [36,37].

This high permeability profile was further corroborated by predicted Caco-2 monolayer transit scores. Critically, neither sesquiterpenoid was classified as a substrate for the P-glycoprotein (P-gp) efflux pump, though both were flagged as potential P-gp inhibitors. This indicates that these compounds might evade active multidrug-resistance efflux mechanisms while modifying substrate transport, which can enhance systemic bioavailabilities [38].

Regarding systemic distribution, the predicted blood-brain barrier (logBB) and central nervous system (logPS) accumulation values fell within acceptable baseline therapeutic parameters [39]. The molecules demonstrated exclusive CYP3A4 selectivity, with no activity against other major CYP enzymes. Predicted CYP3A4 substrate or inhibitory activity points to a potential drug-drug interaction liability requiring experimental confirmation.

Excretion models indicated that predicted total clearance for compounds **1** and **2** was lower than that of doxorubicin and compound 626, suggesting potentially slower elimination; the actual pharmacokinetic significance requires experimental validation [40]. Finally, toxicological screening confirmed a favorable safety profile, showing no predictive liabilities for Ames mutagenicity, cardiotoxicity, hepatotoxicity, or skin sensitization.

In summary, virtual and in vitro evaluations demonstrate that the isolated sesquiterpenoids possess favorable oral absorption profiles and targeted cytotoxic activity against hepatic carcinoma cells, without predicted mutagenic or organ-specific toxic liabilities. This work confirms the thallus tissue of *U. rigida* as a viable ecological niche for isolating *Streptomyces* sp. UL09 and obtaining functional secondary metabolites. Future investigations must focus on validating the safety, maximum tolerated doses, and therapeutic efficacy of these sesquiterpenoid leads *in vivo*.

## 5. CONCLUSION

This study highlights *Ulva rigida*-derived *Streptomyces* sp. UL09 as an alternative bacterial source of insulicolides B and C, which exhibited moderate anti-Gram-positive activity and measurable cytotoxic activity against cancer cells, backed by strong Plk1 docking and favorable ADMET profiles. However, the precise *in vivo* efficacy, mechanistic targets beyond computational modeling, and biosynthetic pathway regulation remain unexplored. Future studies should focus on fermentative yield optimization, *in vivo* pharmacokinetic validation, and detailed mechanistic investigation of these scaffolds for targeted oncological applications.

## 6. ACKNOWLEDGEMENTS

This study was financially supported by the seed grant SRIF-JRG-2569-20 from the Faculty of Science, Silpakorn University, Nakhon Pathom, Thailand.

